# Biases in GWAS – the dog that did not bark

**DOI:** 10.1101/709063

**Authors:** C M Schooling

**Author notes:** **Corresponding author** C M Schooling, PhD 55 West 125.

## Abstract

**Background:** Genome wide association studies (GWAS) of specific diseases are central to scientific discovery. Bias from inevitably recruiting only survivors of genetic make-up and disease specific competing risk has not been comprehensively considered.

**Methods:** We identified sources of bias using directed acyclic graphs, and tested for them in the UK Biobank GWAS by making comparisons across the survival distribution, proxied by age at recruitment.

**Results:** Associations of genetic variants with some diseases depended on their effect on survival. Variants associated with common harmful diseases had weaker or reversed associations with subsequent diseases that shared causes.

**Conclusion:** Genetic studies of diseases that involve surviving other common diseases are open to selection bias that can generate systematic type 2 error. GWAS ignoring such selection bias are most suitable for monogenetic diseases. Genetic effects on age at recruitment may indicate potential bias in disease-specific GWAS and relevance to population health.

## Introduction

Genome wide association studies (GWAS) are a popular and effective way of learning about the causes of diseases. GWAS are particularly valuable because they can give unconfounded estimates of genetic associations, after accounting for population stratification whose importance is emphasized [1]. GWAS are also open to selection bias, the other major source of bias in observation studies [2]. Selection bias, particularly arising due to those missing from a study, is not always intuitively obvious. Over 75 years ago the mathematician Abraham Wald pointed out that findings from a sample of survivors might appear to indicate targets of intervention opposite to the true targets [3]. Specifically, returning fighter planes usually have intact engines, which does not imply the engine is adequately protected, but that only planes with undamaged engines survive, so to improve fighter plane survival the undamaged parts, i.e., engines, require more protection [3]. GWAS are often conducted in samples of survivors, i.e., middle-aged and older people. Genetic randomization on genotype at conception means many years’ worth of selective survival could occur before recruitment [4]. The possibility of bias in GWAS arising from survival on genotype, has been considered and is thought, from simulation, to have little effect on estimates of associations [5] before the age of 75 years [6]. However, many GWAS are of specific conditions. What has not been explicitly considered is the implication for GWAS of considering a particular condition or disease which results in selecting on surviving other causes of death [1]. To avoid such selection bias assessment of the effects of a harmful genotype on a specific condition needs to take into account all other common causes of survival and that condition [7], i.e., competing risk [8]. Here, we explain how and when such selection bias can arise in GWAS, provide empirical evidence together with examples and suggest ways of addressing this bias.

### How selection bias occurs in GWAS

At its simplest the factors determining selection bias in a GWAS are whether the genotype affects survival to recruitment, i.e., selective survival on genotype, and whether other factors cause both the condition of interest and also affect survival to recruitment, i.e., competing risk. For completeness and clarity, we consider all four possible combinations of selective survival on genotype and competing risk, provide the corresponding directed acyclic graphs and give illustrative examples. Directed acyclic graphs allow unambiguous and intuitive presentation of biases as additional open pathways linking the exposure with the outcome. Not fully adjusting for confounders or conditioning on a common factor creates additional biasing pathways [2].

Figure 1a shows no selective survival on genotype and no other causes of the condition of interest, hence no selection bias at all. An example might be a genetic variant for natural hair colour (before going grey), because genetic determinants of natural hair colour are not thought to affect survival and no other factors influence natural hair colour.

**Figure 1:**
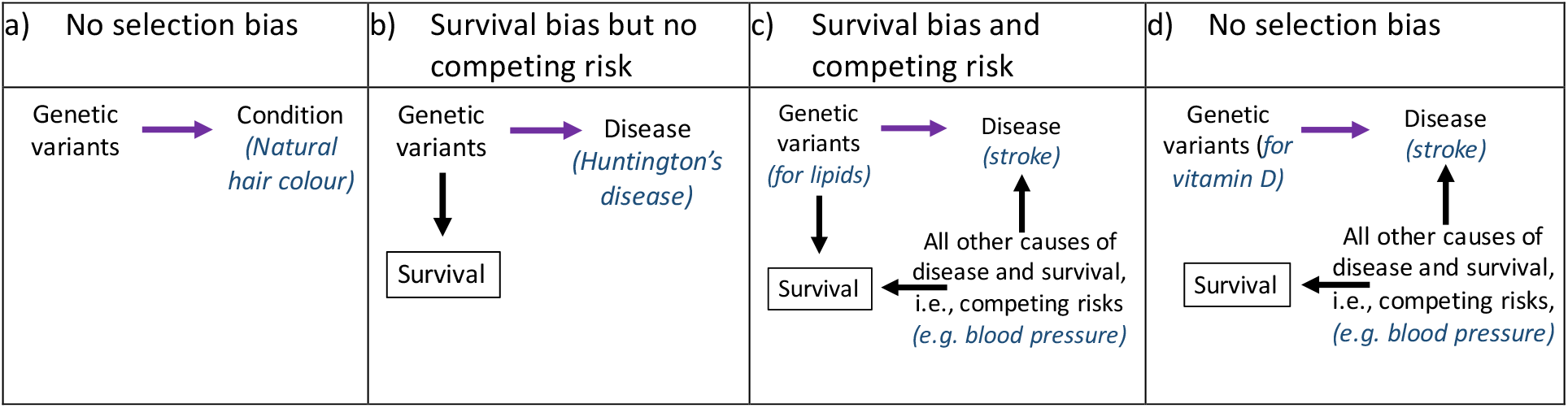
Directed acyclic graphs showing potential sources of selection bias from survival and competing risk. Bias results from any additional open pathway from genetic variant to outcome apart from the association of interest. A link in a pathway may be generated by selecting on a factor, such as survival as here, or by having common causes of survival and disease, i.e., competing risks, that are not fully accounted for in the analysis. If the links together create an alternative pathway from genetic variant to outcome the study will be biased, which can be thought of intuitively as some of the effect coming from or going by a different route so the true effect of genetic variant on outcome cannot be obtained.

Figure 1b shows selective survival on genotype only, with no common causes of survival and the specific condition, and specific effects of genotype on disease. An example might be all genetic determinants of Huntingdon’s disease, because Huntingdon’s disease is a monogenetic disease with specific genetic determinants and no other causes. Estimates may attenuate with increasing age at recruitment because with age the living population available for recruitment and genotyping is increasingly heavily selecting on being alive without Huntingdon’s disease.

Figure 1c shows selective survival on genotype and competing risk from other life-threatening causes of the condition of interest. This situation is particularly likely to occur for conditions that share risk factors with other conditions that cause death before the onset of the condition in question, i.e., for common complex chronic diseases. For example, ischemic heart disease (IHD) and ischemic stroke share several risk factors, such as blood pressure and lipids. However, death from IHD typically occurs in Western populations at younger ages than death from ischemic stroke [9]. As such, genetic associations for stroke are likely at greater risk of selection bias than those for IHD, because regardless of the effect of genotype some people may die from IHD rather than live on to have a stroke, while fewer die from stroke rather than living on to have IHD. This bias may be exacerbated for pleiotropic gene variants that affect common causes of survival and the condition of interest, such as ones associated with obesity.

Figure 1d shows no selective survival on genotype, and hence no selection bias despite common causes of survival and the condition of interest. An example might be genetic variants determining vitamin D on stroke, because vitamin D appears to have no effect on cardiovascular disease or mortality [10] but many other factors affect survival and stroke, such as high blood pressure.

Selection bias arising from missing people who had the exposure (here genotype) and other life-threatening causes of the outcome condition usually biases towards the null, because missing such people from the study dilutes the association in those available for recruitment as they are more likely to be unexposed to a harmful genotype and alive without the disease of interest. Selection bias can also reverse the direction of association. These issues are difficult to spot in genetic studies of complex conditions because few genetic variants have well-established effects. Nevertheless, issues have occasionally been observed in genetic studies but not explained. For example, PCSK9 variants associated with lower low density lipoprotein cholesterol are associated with a lower risk of IHD but not of stroke, although PCSK9 inhibitors protect against stroke [11]. Genetic variants associated with smoking and high blood pressure have been reported as protective against Alzheimer’s disease [12], which seems unlikely.

### Effects of survival on GWAS estimates

To illustrate the effect of survival (on genotype or other causes of the outcome) we chose 6 conditions, 5 non-communicable diseases among the leading causes of years of life lost (chronic airway obstruction, stroke, IHD, colorectal cancer and breast cancer) [13] and diabetes as a contrast because it tends to be a cause not a consequence of life-threatening illness. Table 1 shows a comparison of genetic associations in the UK Biobank for these 6 conditions according to whether the genetic variants were weakly or strongly associated with survival (proxied by age at recruitment). For each condition estimates were on average larger for the genetic variants strongly associated with survival than for the genetic variants weakly associated with survival (Table 1).

**Table 1:**
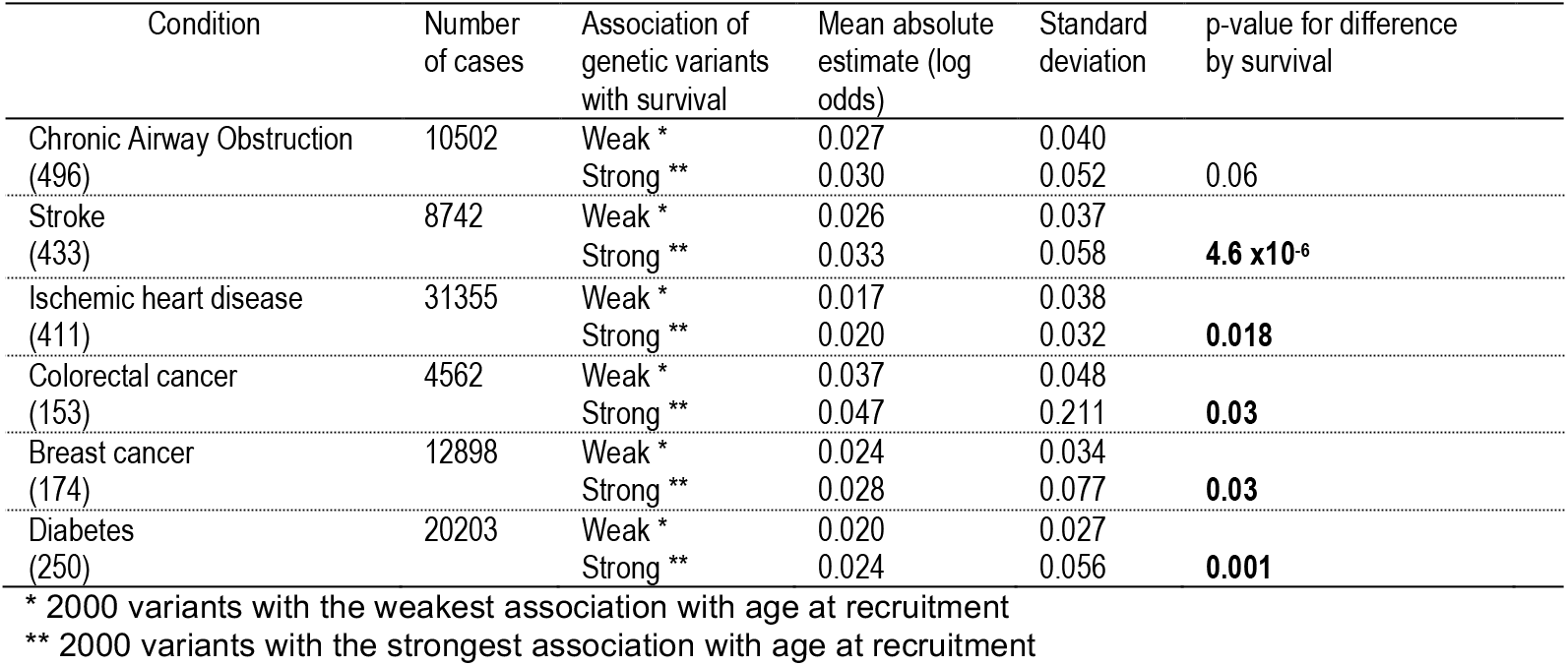
Genetic associations with selected conditions according to whether the genetic variants are very weakly or strongly associated with survival in the UK Biobank

These differences in estimates by genetic associations with survival (Table 1) could be solely due to the genetic variants affecting survival (i.e., survival on genotype) having consistently stronger associations with all conditions or could also be due to survival from other conditions (competing risk) affecting the estimates. To investigate, we compared genetic associations with disease conditions for all these pairs of conditions by the genetic variants’ association with survival. If the conditions are completely independent, then the condition specific genetic associations should have similar differences in magnitude regardless of their association with survival. However, if survival from one condition biases the genetic associations for the other condition then the magnitude of the differences between the two condition specific genetic associations should be more variable for genetic variants associated with survival. Figure 2 shows that for all the 15 unique pairs of the 6 conditions considered, the difference between the estimates was consistently greater and more variable for the genetic variants associated with survival.

**Figure 2:**
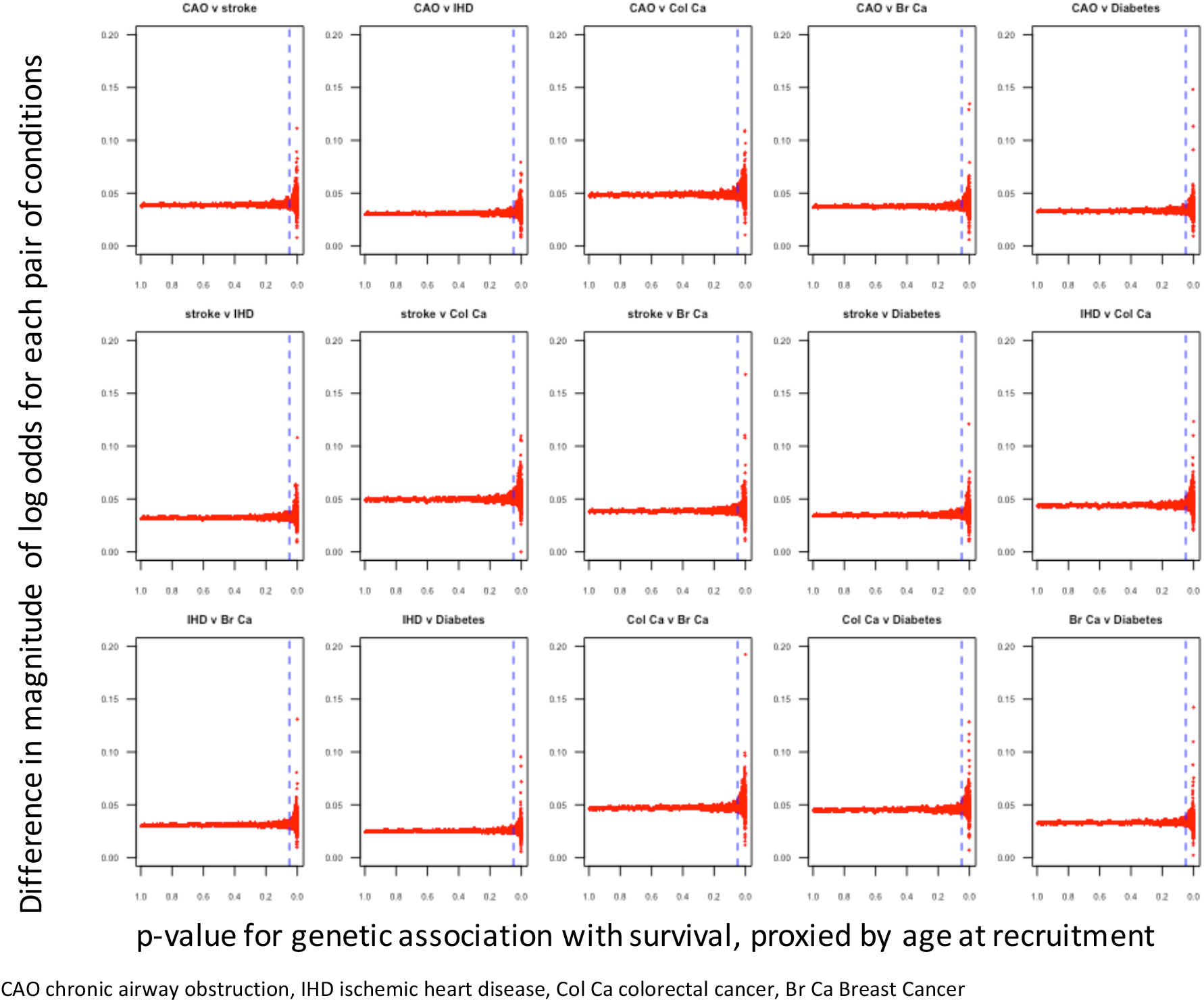
Differences in genetic variant-specific estimates (log odds) for all pairs of the 6 conditions considered according to the effect of the genetic variants on survival, proxied by age at recruitment to the UK Biobank

### Effects of competing risk on GWAS estimates

To identify the effects of selection bias on genetic associations for the SNPs associated with survival, we compared the proportion of genetic estimates, strongly significant for one or other condition, whose magnitude was larger for one condition than another by their association with survival. Given, selection bias usually biases harmful exposures towards the null or even reverses the direction of effect, fewer larger estimates than expected for genetic associations with one condition compared with another for the same genetic variants strongly associated with survival suggests selection bias for the first condition.

Considering the same six conditions (CAO, stroke, IHD, breast cancer, colorectal cancer, and diabetes) we found most evidence that CAO and stroke were missing larger estimates for genetic variants strongly associated with survival compared to genetic variants weakly associated with survival for several of the other conditions, including IHD and colorectal cancer. This observation is consistent with competing risk of IHD and colorectal cancer biasing genetic associations for CAO and stroke towards the null. In contrast, diabetes compared with the other 5 conditions did not appear to be missing any larger estimates for the genetic variants strongly associated with survival than for genetic variants weakly associated with survival (Table 2). This observation is consistent with little competing risk for diabetes. Table 2 also suggests some bias for CAO because of competing risk from diabetes and breast cancer, for IHD from competing risk by colorectal cancer and diabetes, and for breast cancer because of competing risk from colorectal cancer and diabetes.

Selection bias can most simply be eliminated from GWAS by only conducting GWAS on exposures unrelated to survival, for example vitamin D [10]. However, exposures that do not affect survival are of etiological interest but perhaps of less relevance to population health. For exposures that do affect survival only considering conditions not open to competing risk selected from a population where few deaths from related conditions have occurred, for example a GWAS of IHD in middle-aged people rather than stroke in old people, will reduce bias. In fact, selection bias in associations can sometimes be detected by considering how the associations change with age [14]. An association that is most evident in younger people but absent or reversed in older people can be an indicator of selection bias [14]. As such, GWAS routinely stratified by age would be informative. Adjusting for all factors causing survival and the condition of interest is a conceptually simple but a practically infeasible alternative. Conducting a GWAS in a birth cohort without loss to follow-up would avoid bias from selective survival on genotype, because it would eliminate the gap between randomization and recruitment. However, it is not currently very practical and the study would still be open to competing risk for condition specific GWAS. Adjusting genetic associations for the probability of survival, perhaps obtained from genetic associations with age at recruitment, might be helpful but would require additional assumptions to address competing risk. Alternatively, competing risk could perhaps be addressed by a reconceptualization or reclassification of chronic diseases as distinct conditions defined by discrete causes, akin to the way infectious diseases are categorized according to specific causative agents, despite different manifestations.

**Table 2:**
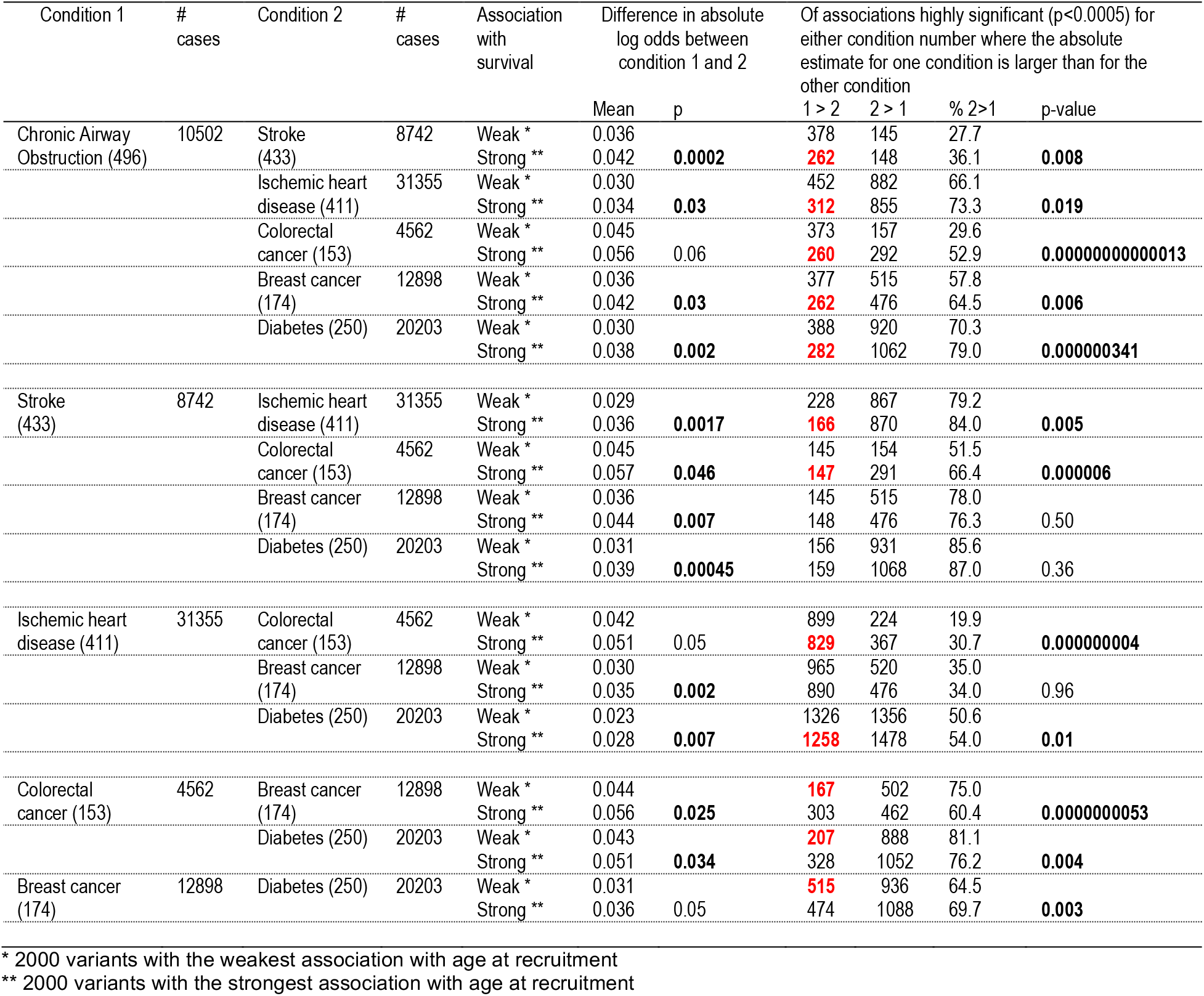
Genetic associations with pairs of conditions by survival in the UK Biobank

### Effects on age at recruitment as a proxy of effects on survival

Given complex chronic diseases may share harmful causes making it difficult to estimate the effect of harmful genotypes on each disease, one possible way forward is to exploit selection bias to obtain effects on survival. Harmful genetic variants inevitably become less common in the population with advancing age, i.e., in surviving older people, meaning associations for harmful exposures will change with age at recruitment into a study. However, genetic variants do not directly cause age at recruitment, so any such association is an indicator of a harmful genetic variant that precluded recruitment, as shown in the directed acyclic graph in Figure 3. This is akin to the practice in studies of aging of comparing the frequency of genetic variants in older and younger people [15], and raises the question as to whether lack of Hardy-Weinberg equilibrium should be taken as an indication of selection bias rather than a reason for exclusion. Here we are proposing to make full use of the study to estimate differences in age of recruitment, which can be interpreted using life tables as years of life lost. Specifically, a genetic variant that results in being a year younger at recruitment means that it gives the same probability of being alive as someone a year older, so its effect can be approximated by the difference in life expectancy for people a year apart in the relevant population.

**Figure 3:**
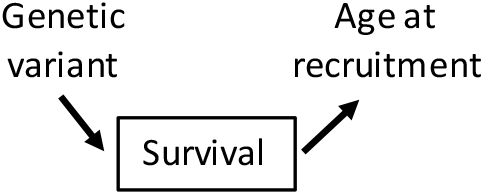
Directed acyclic graph showing the relation of genetic variants to age at recruitment

To illustrate the principle, Table 3 shows associations with age at recruitment to the UK Biobank of some genetic variants with known physiological effects, along with their associations with IHD and stroke also from the UK Biobank. As would be expected, the genetic variants proxying alcohol use and APOE were associated with younger age at recruitment and higher risk of IHD [16, 17]. The genetic variant proxying interleukin-6 was, as expected, associated with lower risk of IHD [18], but had little effect on stroke or age at recruitment. For demonstration several blood pressure genetic variants [19] are shown with consistent associations with IHD and age at recruitment but no association with stroke.

**Table 3:** Associations of genetic variants representing specific exposures with ischemic heart disease, stroke and survival (proxied by age at recruitment) in the UK Biobank

| Exposure | Variant | Effect allele | Ischemic heart disease |  | Ischemic stroke |  | Age at recruitment |  |
| --- | --- | --- | --- | --- | --- | --- | --- | --- |
|  |  |  | OR | 95% CI | OR | 95% CI | beta | 95% CI |
| APOE [23] | rs4420638 | G | 1.09 | 1.07 to 1.12 | 1.09 | 1.05 to 1.14 | -0.10 | -0.14 to -0.05 |
| Alcohol [16] | rs1229984 | C | 1.12 | 1.05 to 1.19 | 1.09 | 0.98 to 1.21 | -0.16 | -0.03 to -0.28 |
| IL6R [18] | rs7529229 | C | 0.97 | 0.95 to 0.99 | 1.01 | 0.98 to 1.04 | 0.02 | -0.02 to 0.06 |
| Blood Pressure [19] | rs3796592 | C | 0.94 | 0.92 to 0.96 | 1.00 | 0.96 to 1.04 | 0.06 | 0.02 to 0.11 |
|  | rs4109837 | T | 1.02 | 1.001 to 1.04 | 1.00 | 0.97 to 1.03 | -0.05 | -0.01 to -0.09 |
|  | rs2306363 | T | 0.97 | 0.95 to 0.99 | 0.98 | 0.95 to 1.02 | 0.05 | 0.005 to 0.09 |

OR odds ratio
CI confidence interval

## Discussion

We have shown theoretically and empirically that GWAS of complex conditions which share causes with common life-threatening conditions that occur earlier in life are open to systematic selection bias (from selection on genotype and surviving competing risk) (Figures 1 and 2) that may reverse the direction of effect, and result in missing genetic associations, i.e., systematic type 2 error in a GWAS (Table 2). However, bias would not be evident for monogenetic diseases that have no other causes and are diagnosed before any deaths have occurred (Figure 1a) where traditional GWAS are likely to be very helpful. We provide a powerful new method for determining whether a genetic variant is likely to be an important target of intervention, even when condition specific GWAS are likely to be biased.

Despite these important findings demonstrating systematic selection bias in condition specific GWAS and providing a method for estimation of effects on a proxy of survival from population representative cross-sectional studies without the need for follow-up, this study is limited in several ways. First, this study does not provide a fool proof method for conducting unbiased GWAS of specific common complex conditions. Methods have recently been developed to tackle the structurally similar problem of obtaining unbiased genetic associations with disease prognosis amongst those with a specific disease [20]. However, the method depends on different factors determining incidence and prognosis, when the same factors may enable survival from a range of chronic conditions [20]. Instead, this study draws attention to the importance of identifying who is unavailable for recruitment from any study sample to avoid selection bias, from selective survival on genotype or competing risk, when assessing potentially causal associations. It also clarifies that condition specific GWAS are most likely to be biased if they are of conditions that share causes with diseases that cause a death at earlier ages, and thereby preclude death from the condition of interest. Second, this study did not consider competing risk after recruitment, because it is more obvious and better understood as it concerns those included in a study not those never available for recruitment. Third, this particular study did not consider all possible conditions open to selection bias because the UK Biobank does not yet have enough cases of all diseases in older people likely to be open to selection bias given the recruitment age was only intended to be from 40 to 69 years [21], the relatively short follow-up to date and the average recruitment age of about 57 years. However, this study is intended to be illustrative and larger similar studies, across a broader age range, could be constructed. Fourth, we assumed that recruitment into the UK Biobank did not vary with age for reasons other than survival. However, it is possible that ill-health also precluded recruitment, which would mean that age of recruitment represents both survival and good health, which does not really affect its interpretation. Fifth, this study is drawing attention to type 2 error, rather than the type 1 error, when is extensively addressed by current methods. Arguably, type 2 is of little importance, however these are systematic type 2 errors. Whether the “missing” associations identified here for disease specific GWAS might be relevant to the small amount of variability explained by such GWAS or the issue of missing heritability could perhaps be considered.

### Conclusion

GWAS of effects of harmful genetic variants on complex chronic diseases are also open to bias from surviving competing risk, which may even reverse the direction of effect. GWAS ignoring this selection bias are most suitable for monogenetic diseases. Techniques to assess genetic effects on complex chronic diseases need to be developed to take account of competing risk before recruitment. Estimating effects of a genetic variant on age at recruitment provides a novel means of obtaining an initial orientation as to whether a genotype and any corresponding exposures are likely to be a useful target for improving population health.

## Methods

### Data sources

We used UK Biobank GWAS as the main source for this paper, because it provides GWAS for many conditions on a common set of participants. The UK Biobank was designed to recruit half a million people aged 40 to 69 years from the UK who were recruited from 2006 to 2010 [21]. Self-reports of health conditions were obtained at baseline with follow-up to all health service encounters and deaths from comprehensive national records.

#### Survival

To proxy survival, we used a GWAS giving associations with age at recruitment to the UK Biobank, because prior death precludes recruitment. The GWAS of 13.7 million variants including up to 361,194 people of white British ancestry adjusted for sex and the first 40 principal components was provided by Neale Lab (http://www.nealelab.is/uk-biobank/). We identified all independent SNPs, i.e., not in linkage disequilibrium (r^2^<0.05), by using the MRBase “clump_data” function.

#### Health Conditions

We choose non-communicable health conditions that are major contributors to years of life lost (YLL) in the UK [13] and currently have enough cases (>4000) in the UK Biobank to generate reliable estimates. Of the 10 leading causes of YLL, we included chronic obstructive pulmonary disease (as CAO), stroke, IHD, colorectal cancer and, breast cancer. We did not include the other 5 leading causes of YLL because they had too few cases (trachea, bronchus and lung cancer, Alzheimer’s disease and other dementias and cirrhosis and other chronic liver diseases) were not available (self-harm) or concerned infectious diseases (lower respiratory infections). We included diabetes as a contrast because it is largely a cause not a consequence of life-threatening illnesses, so it should be less open to survival bias.

To obtain genetic variant specific estimates with these major health conditions we again used GWAS from the UK Biobank here provided by SAIGE which is based on 408,961 white British participants of European ancestry [22]. These genetic associations for 28 million variants were adjusted for sex, birth year and the first four principal components, and used scalable and accurate implementation of generalized mixed models to obtain accurate p-values even when case-control ratios are unbalanced [22].

### Statistical analysis

#### Effects of survival on genotype on GWAS estimates

To assess whether genetic estimates for a condition might be biased by survival, we compared the means of the absolute values of the genetic estimates for each condition across and at the extremes of the survival distribution, where we compared the absolute values of genetic estimates for the 2000 independent SNPs least and most strongly associated with survival using a two-sample t-test (Welch’s unequal variances *t*-test) (Table 1). Estimates unaffected by survival should have a similar magnitude across the survival distribution, while estimates affected by survival may be different for the variants associated with survival.

#### Effects of competing risk on GWAS estimates

To assess how genetic estimates might be affected by surviving other conditions, we assessed for pairs of conditions whether there were differences at the extremes of the survival distribution. First, we plotted the differences against the strength of the association with survival. Given, we had no expectations about the shape of the curve, we simply used an empirical loess plot obtained by ordering the observations by p-value for age of recruitment and then plotting the average difference in small groups (of 71) against the average p-value for same genetic variants. Second, we compared the absolute difference in genetic estimates for the 2000 independent SNPs least and most strongly associated with survival using a two-sample t-test (Welch’s unequal variances *t*-test) (Table 2). Third, we compared the number of times genetic estimates potentially significant (p<0.0005) for at least one of the conditions exceeded the genetic estimate for the other condition (Table 2) for SNPs at both ends of the survival distribution. If genetic estimates for a condition are biased to the null by only observing the survivors of another condition, then the condition affected by such competing risk will have fewer estimates larger than the other condition for the genetic variants associated with survival. We used a chi-squared test to test for the difference. When using the SAIGE UK Biobank GWAS we assumed that effect alleles were used consistently across all conditions.

## Acknowledgements

We would like to thank the groups who provided publicly available GWAS of the UK Biobank. We would also like to thank those who provided encouragement, including Levi Waldron and Eric Tchetgen Tchetgen

